# Alteration of empathy-related behaviors in two strains of mice expressing mutations of the α5 cholinergic nicotinic receptor subunit

**DOI:** 10.64898/2026.01.13.698088

**Authors:** Léa Tochon, Cécile Pagèze, Nadia Henkous, Jean-Louis Guillou, Vincent David

## Abstract

Human genetic association studies have linked a single nucleotide polymorphism (SNP) of the alpha5 subunit of nicotinic acetylcholine receptors (nAChRs) to an increased risk of nicotine dependence, alcohol use disorders (AUDs) and schizophrenia. We used transgenic mice expressing either the SNP rs16969968 (termed α5SNP or α5KI) or a knockout of the *Chrna5* gene (α5KO) to investigate the role of α5-containing nAChRs (α5*nAChRs) in emotion recognition and prosocial, rescuing-like behavioral tasks. We found that α5KO mice are impaired in the recognition of a negative affective state in a familiar peer, and displayed severely altered pro-social, altruistic behaviors, eventually assaulting peers in distress. In contrast, α5KI mice exhibited normal or improved emotion recognition and increased rescuing-like behavior. Importantly, effects of α5 mutations on emotion recognition were modulated by sex. These results demonstrate the critical implication of α5*nAChRs in emotion recognition and prosocial behaviors, revealing, sex-dependent patterns of these social emotional deficits in α5KI and α5KO mice. The current study also supports the view that α5KI and α5KO may provide a valuable preclinical model of Type I (female α5KI mice), and Type II (male α5KO mice) behavioral profiles of AUD.

## 1 INTRODUCTION

Human genome wide studies (GWAS) have reported a link between a single nucleotide polymorphism (SNP rs16969968) of the *CHRNA5* gene encoding for the α5 subunit of nicotinic acetylcholine receptors (nAChRs), and the likelihood of developing nicotine addiction, alcohol use disorder (AUD) and schizophrenia ^1–7^.

The α5 SNP is a frequent missense variation replacing an aspartic acid by an asparagine (D398N), 15% of homozygous carriers showing a two-fold increased risk for smoking ^8,9^. Genetic manipulations of α5 containing nAChRs (α5*nAChRs) in rodents have linked a null mutation or SNP of *Chrna5* with differences in alcohol self-administration and tolerance ^7,10–12^. Transgenic rats expressing the α5SNP through a genetic knock-in (α5KI) procedure, consume more alcohol, and exhibit increased relapse to alcohol seeking after abstinence ^13^. Consistently, we have demonstrated that both α5KI and α5KO mutant mice showed alcohol over-consumption for highly concentrated solutions, but display opposite anxiety-related behavior (hyper vs hypo-anxious) and behavioral control (not impulsive vs impulsive-like, respectively) ^7^. We pointed out that these opposite phenotypes strongly evoke the two opposite human subtypes of AUD initially referred to as Type I and Type II based on different characteristics such as, personality traits, biological sex and age of onset ^14,15^. Cloninger’s *Type I* (‘*Avoidance alcoholism*’) is attributed to middle-aged individuals who are prone to anxiety and who are thought to indulge alcohol in a ‘self-medicating’ way to reduce stress and cope with their symptoms ^14^. At the other end of the continuum, Cloninger’s *Type II* (‘*Sensation-seeking alcoholism*’), which has a male predominance, is characterized by impulsivity and sensation seeking behavioral traits accompanied by antisocial behavior and displays of aggression; for this subtype, alcohol is suggested to be used to enhance their sense of thrill ^14,16^.

In light of our previous data showing behavioral similarities between α5KI and α5KO mutant mice, the Cloninger’s avoidant (Type I) and sensation-seeking (Type II) AUD subtypes respectively, we raise the hypothesis that α5 nicotinic mutants may be a relevant basis for preclinically modelling AUD subtypes ^7^. However, one remaining issue about the relevance of these mouse models concerns their social-emotional profile. AUD is associated with difficulties in emotion processing and social cognition, including deficits in empathy-related processes like facial emotion recognition ^17–26^. Empathy in its broad sense can be defined as an induction process that reflects the cognitive ability to perceive the emotional states of others and the affective tendency to experience emotions of others, which can be coupled with the motivation to care for their well-being and to help by acting prosocially (i.e acting to benefit another) ^27,28^. The difficulty to respond appropriately to another’s emotions can compromise interpersonal relationships, thereby contributing to the onset and/or the sustainment of alcohol-related problems but also interfere with recovery ^21,29^. However, the types of social-emotional impairment seem to differ across AUD subpopulations as observed in Cloninger’s Type I and II. The first are described as ‘eager to help others’ and ‘sensitive to social cues’ but seemed to misinterpret facial expressions of sadness and contempt ^14,30^. At the opposite, the latter were found to overinterpret facial expressions of anger and disgust, showed a preponderance of anger mimicry and aggressive behaviors ^14,31^ which is associated with deficient affective empathy ^32^ and is congruent with their high rate of comorbidity with antisocial personality disorder ^14,31^. There is therefore a crucial need for better characterization of emotional and social cognitive processes to understand the different motivations leading to alcohol abuse and to develop suitable interventions taking into account patients’ socio-emotional skills ^33^.

In the past years, rodents have been increasingly used to study social capabilities considered as empathy-related features ^34–38^. New experimental protocols have been developed to assess their emotion recognition abilities ^39–43^ and prosocial motivation to help ^44–49^. Moreover, α5-nAChRs are highly expressed in key brain structures involved in social cognition and social-emotional processing such as the amygdala, prefrontal cortex ^50,51^, subiculum, ventral hippocampus ^52,53^, olfactory bulb and hypothalamus ^54,55^. We thus investigated the functional impact of either the α5SNP or a null mutation of *Chrna5* on emotion recognition abilities and prosocial, altruistic behaviors in mice. We hypothesized that either α5KO and/or α5KI would perform differentially from WT mice on these social tasks, thus demonstrating the implication of α5-nAChRs in socio-emotional behaviors and potentially furthering the analogy between these mutant strains and AUD subtypes behavioral characteristics ^14^.

## 4 MATERIALS & METHODS

### 4.1 Animals and housing

We used two transgenic mouse strains expressing different mutations of the *Chrna5* gene encoding for the α5 subunit of nAChRs, on a C57BL/6J background. The α5KO mice carry a null mutation of *Chrna5,* while the α5SNP mice (referred to as α5KI hereafter) express a single nucleotide polymorphic (SNP) variant (rs16969968) defined as a substitution of the aspartate 398 by an asparagine (D398N), corresponding to the most frequent α5-SNP found in the Western Population ^56^. They were initially provided by our collaborators at the Pasteur Institute (Paris), and were bred in the animal facility of the lab for 8 generations with a cross-breeding with heterozygous mice from the Pasteur Institute at the 4^th^ generation. Wild-type mice were obtained from the same litter as both α5 nicotinic mutants. Genotype identifications were conducted at 7 post-natal days (P7) through PCR analysis of tail samples by the genotyping platform of the Neurocentre Magendie in Bordeaux. For the experiments, we used mice aged of 20 to 24 weeks. The mice were housed in collective cages (4 to 8 per cage) with access to water and food *ad libitum* in a temperature-controlled room (22 ± 1°C). The lighting of the cages followed a 12-hour day/night cycle in such a way to allow testing to occur from 7:00 AM to 07:00 PM during the mice’s day time, following approval of the Local Ethics Committee and in agreement with the CEE Directive 86/609/EEC.

### 4.2 Group repartition

Each mouse underwent two behavioral tasks: the affective state discrimination task (ASDT), and the restrainer tube test (RTT) 4 weeks later. Both male and female mutants (α5KO: n=8 ♂ and 5 ♀; α5KI: n=7 ♂ and 9 ♀) as well as male and female WT littermate mice (n=9 ♂ and 9 ♀) were included in these experiments.

### 4.3 The Affective State Discrimination Task (ASDT)

We used an adaptation of the ASDT protocol described by Scheggia and colleagues (2019), which consist in determining the preferential approach tendency in observer mice when exposed to two unfamiliar demonstrator mice in different affective states ^39,43^. This aimed at testing the capability of mice to discriminate between a neutral demonstrator and a stressed demonstrator, as part of an emotion recognition task. Briefly, the observer was introduced in a cage (29 x 35 x 19 cm) containing two demonstrator mice placed in transparent plastic cylinders (H:21 cm; D:10 cm) with perforated holes at the bottom to allow for breathing and sensory interaction with the environment. A metal separator (2mm x 21 cm x 18 cm) was placed in the middle of the cage in order to mark off the two areas of interest corresponding to each demonstrator and blocking their reciprocal view of each other. Demonstrators were WT mice and were sex-matched to the observer. The entire testing arena was wiped down with warm water and the litter was changed between each subject to eliminate any odors left by the mice. At the end of each testing day, the entire apparatus was wiped down with Ethanol (70%).

#### Habituation

All the mice (demonstrators and observers) were handled for 5 minutes a day over 3 days to become familiar with the experimenter ^57^. They were then habituated to the testing zone for 10 minutes a day over another 3 days. For the test zone habituation, observer mice were gently placed in the ASDT arena and allowed to freely explore in the absence of the demonstrator mice inside each cylinder (Figure 1, top). During the habituation, the mice’s movements were tracked with a tracking Software (Anymaze, Stoelting). After the observer mouse’s habituation trial ended, its allocated pair of demonstrator mice was subsequently gently placed inside their corresponding cylinder for 10 minutes.

**Figure 1.**
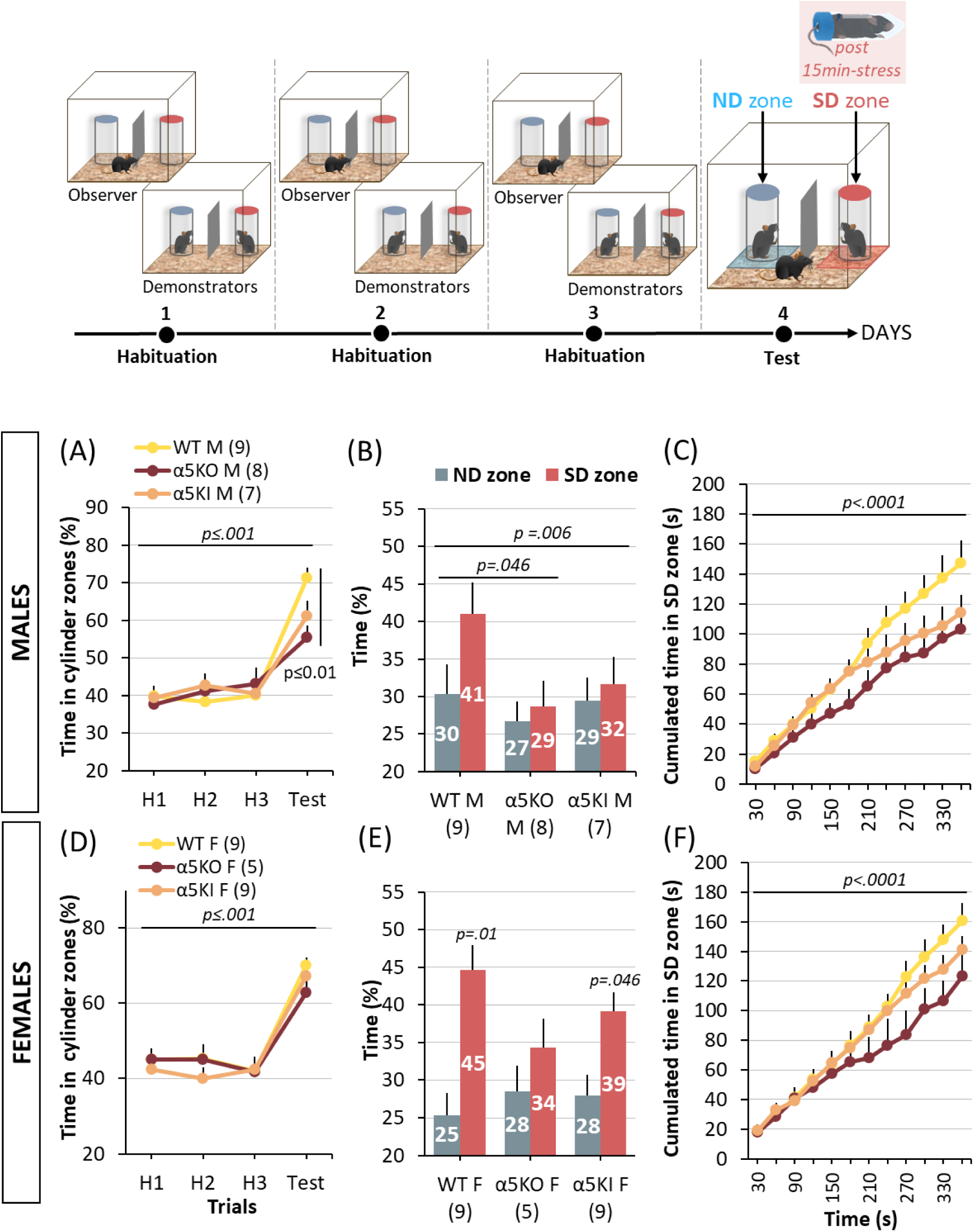
Genotype and Sex-dependent emotion recognition deficits in α5 nicotinic mutant mice Top: Time course and protocol of the Affective State Discrimination Task (ASDT). Proportion of time spent in the cylinder zones surrounding the demonstrators during the three habituation trials and the test day across α5 nicotinic mutants and controls in males **(A)** and females (**D).** Proportion of time spent in the neutral demonstrator (ND) zone (Blue) and stress demonstrator (SD) zone (Red) respectively during the test across α5 mutants and controls in males **(B)** and females **(E).** Proportion of time spent in the SD zone during the test across time (30 second segments) for α5 mutants and controls in males **(C)** and females **(F)**. *F, females; M, males*. p≤0.001: trial/time effect (RM-ANOVA); p≤0.01: genotype x trial/time interaction (RM-ANOVA), p≤0.01: genotype effect (RM-ANOVA); p≤0.01: α5KO vs WT (Bonferroni/Dunn post-hoc); p≤0.05; p≤0.01: SD vs ND (t-test; paired). p≤0.001. ND: neutral demonstrator; SD: stressed demonstrator

#### Testing

For the purpose of creating different affective states in mice, the demonstrators were divided into 2 groups: *Neutral* and *Stress*. *Stress* demonstrators (SD) underwent a restraint procedure for 15 minutes prior to the beginning of the test which consisted in constraining them inside a 50mL custom-made tube (*Falcon©*) perforated at the end to allow for breathing. After 15 minutes of restraint, the stressed demonstrator was immediately placed in one of the two cylinders which was marked by a red-lid for identification. *Neutral* demonstrators (ND) were not further handled in a way to avoid altering their emotional state and were directly transferred from their home cage to the other cylinder which was marked with a blue lid. For each test trial, ND and SD mice were randomly allocated either to the right or left cylinder relative to the central separator, during both habituation and testing trials, to avoid the confounding variable of spatial preference. The observer was then gently introduced inside the arena and its exploration was recorded for 10 minutes using Anymaze. *Demonstrator zones (SD* and *ND zones)* were outlined using the tracking software and consisted in a square perimeter of 2 cm around and including the cylinders. Parameters of interest such as: time spent and number of entries inside the zone were automatically recorded. In addition, sniffing of the demonstrators were scored *a posteriori* blindly using the off-line recordings.

##### ASDT parameters

###### Parameters recorded were time spent around each demonstrator zone (SD and ND zones)

Quantification of sniffing behavior towards SD vs ND was used to establish a preference score defined as SD sniffing – ND sniffing. A sniffing event was operationalized by the occurrence of a nose contact of the observer towards the cylinder containing a demonstrator. We then categorized each group into discriminators by using a cut-off score of 10 sniffings (Pref. Score >10; high discriminators) versus non-discriminators (Pref. Score <10).

### 4.4 The Restrainer Tube Test (RTT)

The Restrainer Tube Test (RTT) aimed at exploring pro-social rescuing-like behavior through the testing of social release. We based our testing procedure on the ones used by Bartal et al. (2011) and Fontes-Dutra et al. (2019) in their studies measuring the willingness of rats to free their cagemate trapped in a tube and its adaptation to mice by Ueno et al. (2019). The test was carried out over 5 days following 4 different phases shown in Figure 3: *Self-freeing training* (Day 1 & 2), *Chocolate versus Empty tube* (Day 3), *Cagemate versus Empty tube* (Day 4) and a competition trial: *Cagemate versus Chocolate tube* (Day 5). In contrast to protocols established for rats in which trap devices may not be stressful enough ^58^, the adaptation to mice using a restraint tube ensure a significant distress ^59^. Moreover, we develop a short procedure with only 2 days of exposure to entrapment (for both subject and cagemate mice), this period of time is not sufficient for the mice to develop stress habituation ^60^.

#### Self-freeing training

The mouse was placed in the restraint tube (50mL plastic tube, Falcon©) closed off with a paper door at the extremity facing the head of the animal inside the testing cage, the mouse therefore learned to free itself from the inside of the tube by chewing through the paper door. The training was done three times a day over two days (3 trials on Day 1 + 3 trials on Day 2; Figure 3, top). All the mice successfully learned how to open the tube from the inside after 2 days of training.

#### Chocolate Opening Training

The mouse was trained to open the paper door on a restraint tube from the outside to access white chocolate chips (3 x 25 mg chips). In order to motivate the animals for the task, they were food-deprived during the 12 hours preceding the training. The mouse was placed inside the testing cage with an empty and a chocolate containing tube closed off with a paper lid and was left to roam freely until it had opened the chocolate tube and eaten the chocolate chips. This training was done 3 times over one day (3 trials on Day 3; Figure 3).

#### Cagemate Freeing Training

Each test mouse was assigned a sex-matched ‘*trapped cagemate’* consisting in a heterozygous (+/-) littermate that was trapped inside a paper sealed tube at its rear so that the trapped mouse could not free itself. The test mouse was placed in the cage with the cagemate-containing tube as well as an empty paper-sealed tube. In order to free its cagemate, the test mouse needed to chew through the paper door from the outside. The session ended 5 minutes after the mouse had successfully freed its cagemate, or after a maximum of 90 minutes had elapsed. The training was repeated twice over one day (2 trials on Day 4; Figure 3).

#### Chocolate versus Cagemate trial

The final trial of the test consisted in placing the mouse in the testing arena in the presence of a chocolate-containing tube (3x 25 mg white chocolate chips) and a cagemate-containing tube, both closed off with a paper lid (1 trial on Day 5; Figure 3).

For all trials, the latency to open the tubes as well as the order of opening of the tubes was recorded. Additionally, the number of white chocolate chips eaten by the test mouse was counted to account for sharing-like behavior. A digital camera was placed above the testing apparatus to record the Chocolate versus Cagemate Trial with a behavioral tracking system (Anymaze, Stoelting). *Tube zones* were outlined with the software which consisted in a rectangular perimeter of 1 cm around each tube, and accordingly parameters such as time spent in the tube zones and number of entries inside each tube zone was recorded.

##### Score for empathy-like behavior

To assess the empathy-like profile of nicotinic mutants as compared to WT controls, a score was calculated according to two factors: namely, *Cagemate Prioritization (CP)* which allocated a score between −3 and 3 to the mice depending on whether the cagemate tube (+1), the chocolate tube (−1) or the empty tube (−1) was opened first during trials *cagemate versus empty* and *cagemate versus chocolate*. The second factor was *social-contact positivity (SCP)* and allocated a score between −3 and 3 to mice depending on whether interacting (+1), disinterested (0) or aggressive (−1) behavior was displayed towards the cagemate after the test. The interaction type was categorized in real-time during the test by the experimenter: *Interactive* behavior was attributed when a minimum of 10 sniffings where observed, on the contrary when less than 10 sniffings were observed and the mice spent most of the interaction time at opposite ends of the cage, the interaction was considered as *disinterested*. *Aggressive* behavior was considered when 2 or more aggressive events were recorded between the test mouse and the cagemate (chasings and/or biting). In the latter case, the session was immediately stopped. Score formula:

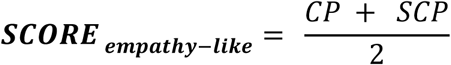

### 4.5 Statistical Analysis

Distribution of samples was tested using Kolmogorov-Smirnov test. Behavioral performance data following normal distribution were analyzed using an ANOVA with one or multiple factors (genotype/RM, repeated measures), followed by appropriate post-hoc tests (Bonferonni/Dunn) to compare the different groups. Frequencies of specific behavioral responses across groups in the ASDT were compared using Chi Square (*X*^2^) tests. Empathy-like scores in RTT were compared either using the non-parametric Kruskal-Wallis test (for comparison between the 3 genotypes, within and without sex); or using the non-parametric Mann-Whitney U test (for direct comparison between 2 genotypes, within and without sex groups).

## 2 RESULTS

### 2.1 Emotion discrimination in α5KO and α5KI mice

We first assessed the ability of α5KO and α5SNP (=α5KI) transgenic mouse strains to discriminate between two unfamiliar mice (called demonstrators) in different affective states, i.e. either a state of psychological stress induced by 15 minutes of constriction inside a tube (SD), or a neutral state (ND). We found both strains to be sensitive to the presence of demonstrator mice inside the cylinders, as revealed by the time spent in the demonstrator zones during the test trial as compared to the three habituation trials in all groups (M + F; trial effect: F_(3,123)_=116.46, *p<0.001;* sex effect: F_(1,41)_=2.81, *p=0.10;* Figure 1A, D). In WT mice, time spent in the SD zone relative to the ND zone was higher similarly for both sexes (t(17)=3.14, *p=0.006;* Figure 1B, E). In contrast, α5KO mice (both male and female) appeared severely impaired in this task spending an equivalent amount of time in each zone (t(8)= p=0.33; Figure 1B, E). α5KI exhibited an intermediate performance. Interestingly however, while male α5KI appeared severely impaired in this task (t(8)=, *p=0.70;* Figure 1B), their female counterparts were still able to discriminate between SD and ND demonstrators (t(8)=2.36, *p=0.046;* Figure 1E).

The recording of the cumulated time spent in the SD zone throughout the session enabled us to visualize kinetics of the interest exhibited towards the stress demonstrator over the 6-minute duration of the trial (Figure 1C, F). In WT mice, time spent in the SD zone remained constant throughout the 6-minute session, indicating a sustained interest for the SD. However, both male and female α5KO explored less the SD zone from 210 to 360 sec than WT mice, whereas α5KI exhibited an intermediate performance (effect of genotype effect, *F_(2,44)_=4.73, p=0.02*; effect of time, *F_(1.89,83.58)_=323.7, p<.0001*; genotype x time interaction (*F_(22,484)_= 4.73, p<0.0001*; Post-hoc Bonferroni/Dunn WT vs α5KO: *p=0.016,* WT vs α5KI: *p=0.33*). There was no effect of sex on this parameter.

Quantification of sniffing behavior towards SD vs ND provided further information about affective state discrimination in both nicotinic mutant strains. While WT observer mice showed a clear preference toward SDs, particularly females (t(8)=9.57, *p*<0.0001), this behavior was totally suppressed in most of null mutants (males: t(7)=.42, p=0.68; females: t(4)=0.38, p=0.72) (Fig. 2A,C. for examples of these behaviors see Video 1 and 2).

**Figure 2.**
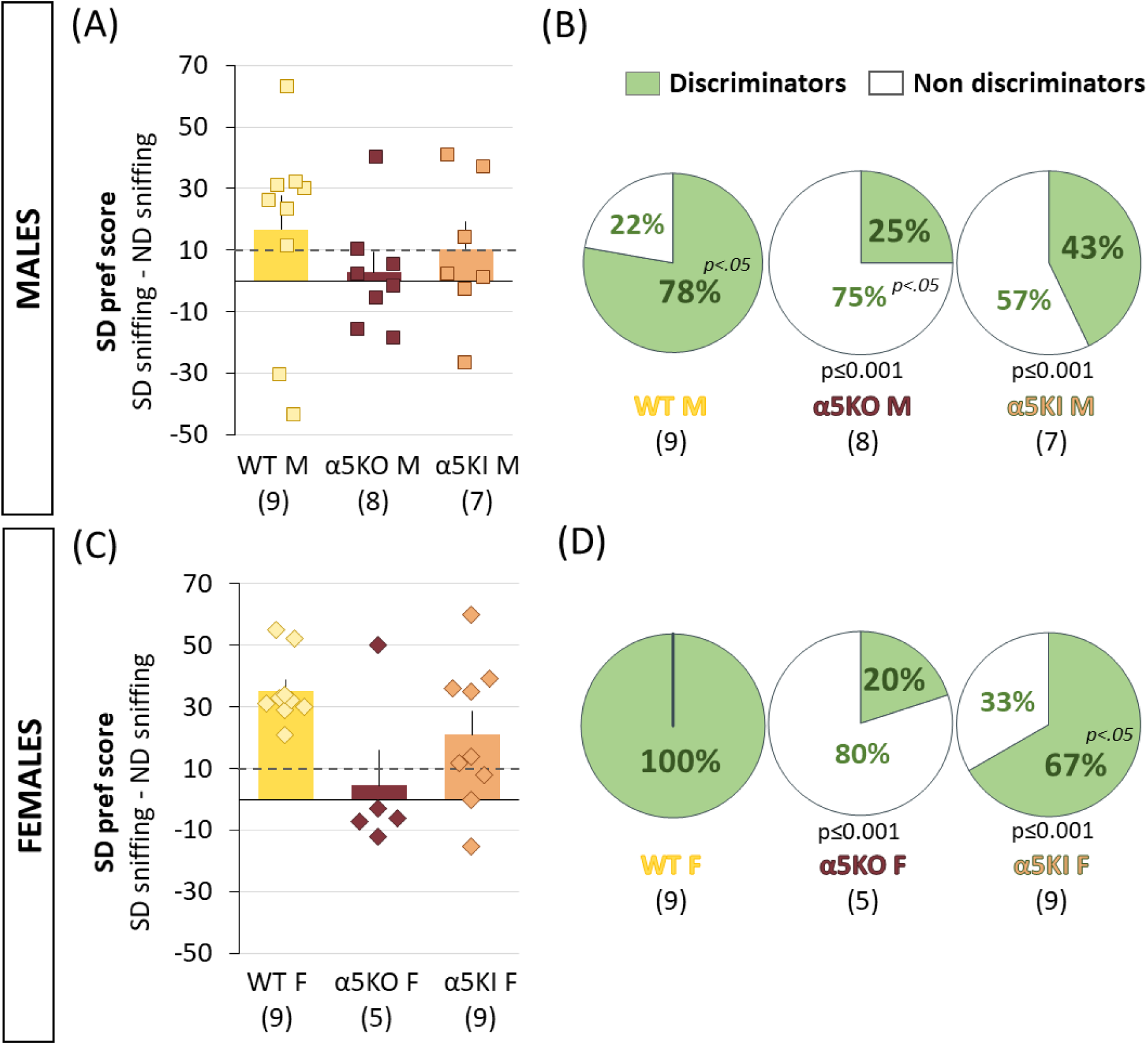
Impairment of exploratory behavior towards a stressed demonstrator in α5 nicotinic mutants. SD preference score corresponding to the number of sniffings inside the ND zone subtracted to the number of sniffing inside the SD zone across α5 nicotinic mutants and controls in males **(A)** and females **(C).** A cut-off score is set at 10 to differentiate between discriminators (Preference score > 10) and non-discriminators (Preference score < 10) whose relative proportion for each genotype is shown in **(B)** for males and in **(D)** for females. *F, females; M, males.* p≤0.001: genotype x profil interaction (X^2^ vs WT); p≤0.05: discriminators vs non discriminators (X^2^).

Counting the number of discriminator (Pref. Score >10) vs non-discriminator (Pref. Score <10) mice within each group revealed further interesting information. First, although both male and females WT mice all performed well on this task, female were unanimous affective discriminators when 22% of male WT were non-discriminators (M vs F: *X*^2^=40.99, n=18, *p<0.0001;* Figure 2B vs 2D). The repartition of these two profiles (discriminators vs non-discriminators) was strikingly inverted in null mutant male and female mice. Finally, KI mice again showed an intermediate distribution (WT vs KO: *X*^2^ =4.2, *p=0.0031*; WT vs KI: *X*^2^ =3.7, *p=0.0031*; Fig. 2B, D).

### 2.2 α5KO and α5KI Mice Display Similar Cagemate Freeing Behavior

We then aimed at investigating the altruistic behavior of α5 nicotinic mutants using the Restrainer Tube Task (RTT). Over the first two days, the mice underwent training to learn how to open a paper-sealed tube first on the inside while trapped inside the tube, and from the outside when they were confronted with a tube containing white chocolate chips after being food deprived for 12 hours (Figure 3, top). All mice, regardless of genotype, efficiently learned to open the paper sealed tube from the inside as witnessed by the decrease in their latency to self-freeing (M + F, mean latency on trial 1 vs trial 6: 335±47.8 vs 70±9 sec); trial effect: F_(5,205)_=13.52, *p<0.0001*). Notably, females were quicker to free themselves compared to males (mean latency on 6 trials in M vs F: 91±8.5 vs 200±21.7 sec; sex effect: F_(1,41)_=21.26, *p<0.0001;* trial x sex interaction: F_(5,205)_=2.89, *p=0.015).* During the Chocolate versus Empty phase, all the mice regardless of genotype or sex were quicker to open the chocolate containing tube on the third trial compared to the first (mean latency on trial 7 vs trial 9: 1116±97.6 vs 367±65.6 sec; trial effect: F_(2,82)_=35.39, *p<*0.0001; Bonferroni/Dunn post-hoc: *p<*0.0001), indicating that they all efficiently learnt how to open a paper sealed chocolate-containing tube from the outside. There was no effect of genotype or sex on the opening latency for the chocolate containing tube.

**Figure 3.**
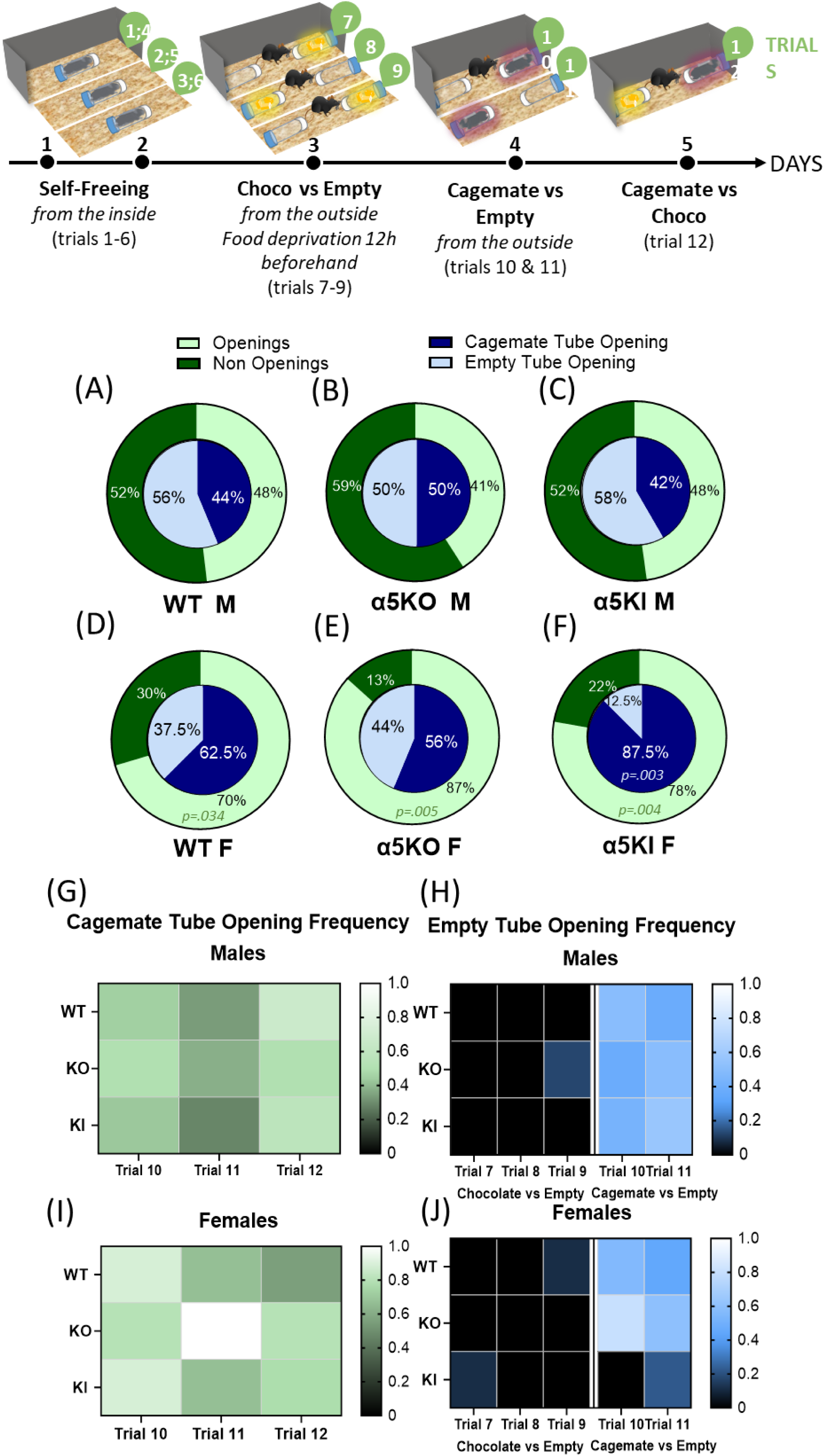
Female a5KI mice exhibit increased cagemate freeing. Top: Time course and detailed protocol of the Restrainer Tube Task (RTT) spread over 5 consecutive days (black dots) consisting in a total of 12 trials (green bubbles). **A-F**: Pie charts of the proportions of cagemate openings, *“Cagemate Openings”* were considered as such when the test mouse opened the cagemate-containing tube more than once over the three trials (light green) and “no cagemate openings” considered as such when they fail to open the cagemate tube over the three trials (dark green) along with the proportion of cagemate tube openings (dark blue) versus empty tube openings (light blue) in the trials containing a cagemate tube and the trials containing empty tubes respectively for male WT, α5KO and α5KI mice **(A,B** and **C)** and female WT, α5KO and α5KI mice **(D, E, F). G:** Frequency chart for cagemate tube openings across the three trials in which this outcome is possible (presence of a trapped cagemate among the presented tubes) in males and in females **(I)**, lighter gradient corresponds to a higher frequency**. (H)** Frequency chart for the empty tube opening frequency during the trials in which this outcome is possible, this include trials in which a chocolate and an empty tube are present in the arena (Trial 7-9) and trials when a cagemate and an empty tube are present in the arena (Trial 10-11) in male and females **(J),** lighter gradients correspond to higher frequency. *CM, cagemate; F, females; M, males*.p<0.05, p<0.01 (X^2^).

Concerning the cagemate tube opening behavior (Cagemate versus Empty and Cagemate versus Chocolate), a distinction of opening profiles was done based on whether a minimum of two successful cagemate freeing events occurred out of the total three trials (Figure 3, see Video 3). WT, α5KO and α5KI mice displayed similar opening profiles, but females were freeing their trapped cagemate more than their males counterparts did (χ2(1,18)=40.99 p<0.0001, Figure 3A vs 3D; χ2(1,12)=3.03 p=0.08, Figure 3B vs 3E; Figure 3C vs 3F; χ2(1,12)=86.92 p<0.001). All groups opened both empty and cagemate tubes with a similar frequency (Figure 3 G-J), but females more specifically α5KI opened more cagemate tubes than empty ones (respectively: 88% versus 12%; χ2(1,18)=16.39, p<0.0001; Figure 3I,J). This result indicates that the tube opening behavior of α5KI females was guided by the presence of a cagemate inside the tube.

### 2.3 α5KI Mice but not α5KO mice display increased altruistic behaviors

During the competition phase between one tube containing a chocolate chip and another one containing a cagemate tube, most of the mice, regardless of their group, spent more time in the surrounding of a chocolate**-**containing tube when compared to the cagemate tube (tube effect, M+F: F_(1,44)=_27.67, *p<0.0001; M:* F_(1,21)=_29.91, *p<0.0001*; F: F_(1,20)=_7.73, *p=0.011;* Figure 4A, B). However, there was a clear preference of α5KI mice toward the cagemate tube, as compared to both WT and null mutants (expressed as the number of entries inside the cagemate tube zone subtracted to the entries inside the chocolate tube zone). This is supported by a main effect of genotype on this variable (M+F: F_(2,41)_=3.42, *p=0.042*; Bonferroni/Dunn post-hoc, α5KI vs WT: *p=*0.011; Figure 4C), indicating an increase in prosocial behavior in α5KI mice.

**Figure 4.**
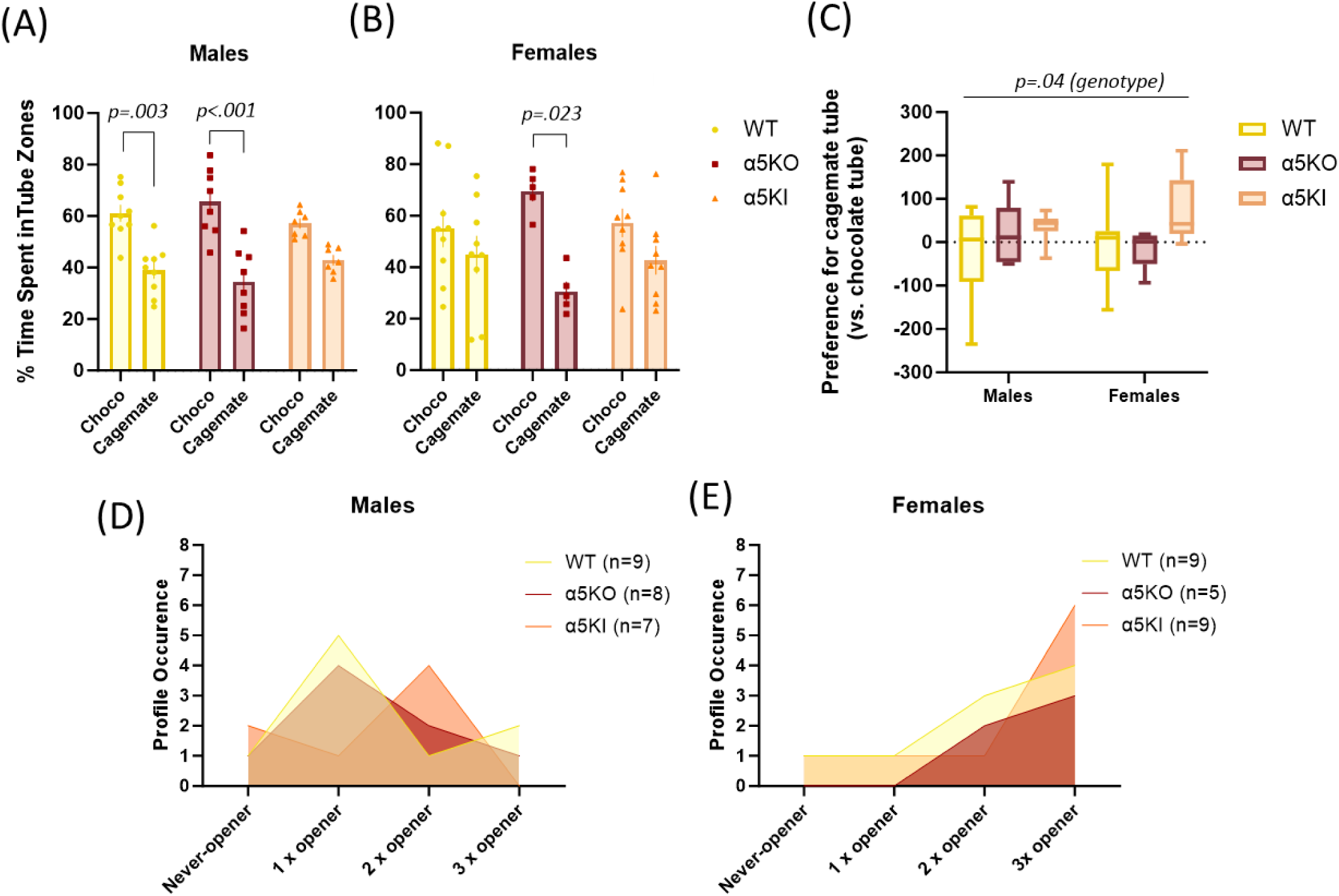
Mice prefer Chocolate over Cagemate containing tubes, but α5KI mice show increased pro-social behavior. Proportion of time spent in each tube zone during the Cagemate versus Chocolate trial for males **(A)** and females **(B).** Cagemate vs Tube Preference Score corresponding to the number of entries inside the cagemate tube zone subtracted to the number of entries inside the chocolate tube zone across male and female groups of α5KO, α5KI and WT **(C).** p≤0.05; p≤0.01; p≤0.001: time in the Choco vs CM tube zone (t-test; paired); *p<0.05*: genotype effect (RM-ANOVA). **D, E.** Plot of the occurrence of different profiles relating to cagemate opening across each group in males (left) and females (right).

### 2.4 α5KO and α5KI Mice Display Opposite Post Freeing Interactions

To further characterize the prosocial profile of both mutant strains, we scored social interactions immediately following the exit of the cagemate from the tube at the end of the test (Figure 5). The results revealed a striking opposition between α5KO and α5KI mice, with a strong effect of sex. WT mice of both sexes (but more so females) engage mainly in positive interactions after cagemate freeing. In contrast, male α5KO exert higher levels of aggression towards their cagemate as compared to either WT or α5KI males (genotype effect: F_(2,21)_=5.27, *p=0.013*; Bonferroni/Dunn post-hoc, α5KO vs WT: *p=0.0079*, α5KO vs α5KI: *p=0.014*; Figure 5A left and Video 4). This aggressive response is not observed in α5KO females, which are predominantly disinterested by the cagemate (genotype effect: F_(2,20)=_3.68, *p=0.043;* Bonferroni/Dunn post-hoc, α5KI vs α5KO: *p=0.014,* α5KO vs WT: *p=0.051 ns;* Figure 5A right). Conversely, female α5KI express a more interactive, WT-like behavior (genotype effect: F_(2,20)=_3.58, *p=0.046;* Bonferroni/Dunn post-hoc, α5KI vs α5KO: *p=0.014,* α5KI vs WT: *p=0.29 ns;* Figure 5A right).

**Figure 5.**
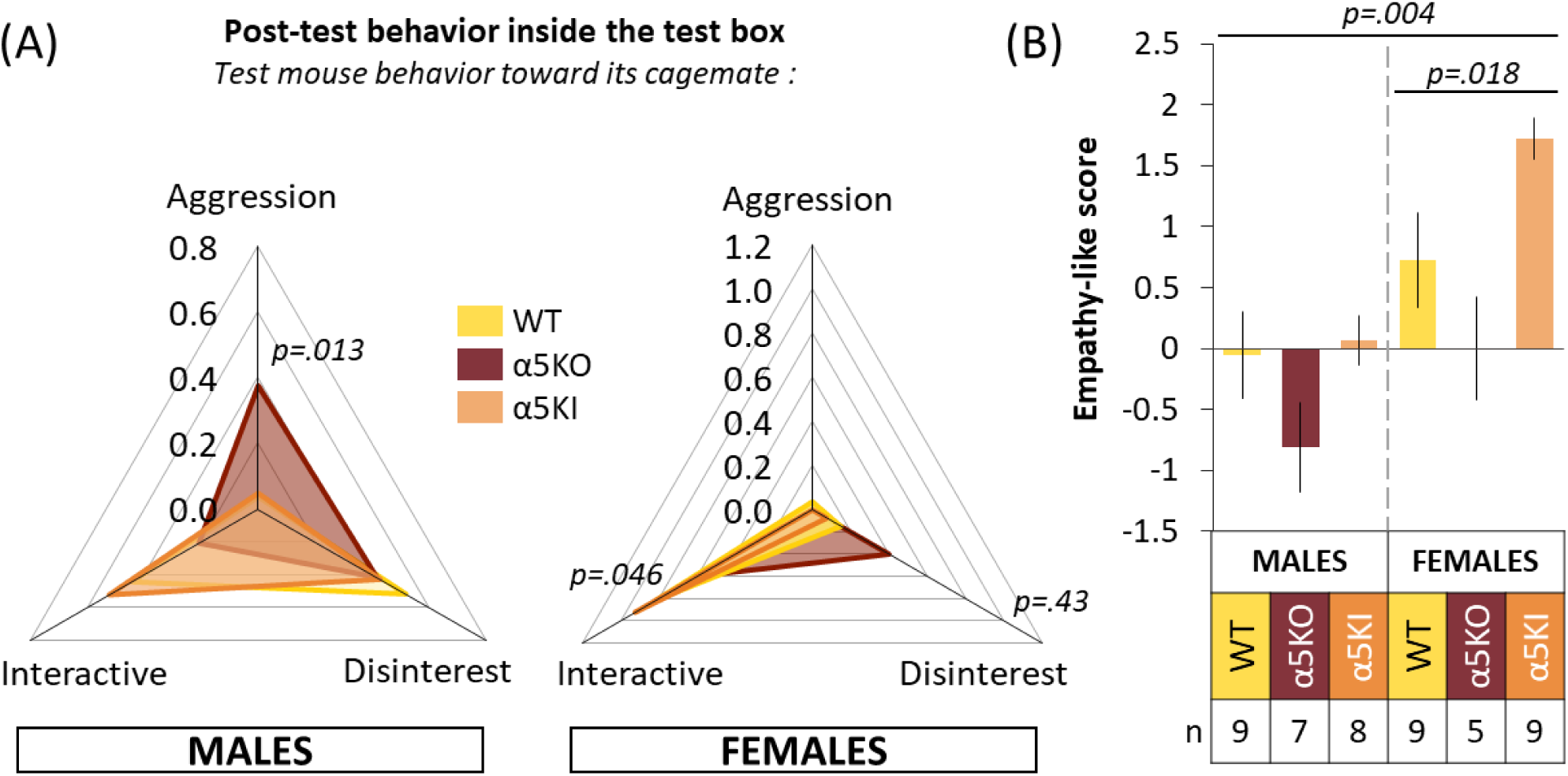
α5 mutants display opposite interaction profiles with the freed cagemate. **(A)** Representation of the frequency of aggression, interactive or disinterested interaction for male WT, α5KI and α5KO mice (left) and female WT, α5KI and α5KO mice (right). **(B)** Empathy-like score for empathy across male and female nicotinic mutants and controls. The empathy-like score is calculated according to two factors: *Cagemate prioritization* (depending on which tube is opened first between cagemate, chocolate and empty) and *social-contact positivity* according to which type of interaction the mice partake in with the cagemate following the test (Aggression, interactive or disinterested).*p<0.05: Genotype effect (ANOVA); p<0.05: Genotype effect (Kruskal-Wallis test); p<0.05: Genotype effect in females (Kruskal-Wallis test)*.

Scores of empathy-related behavior based on both cagemate prioritization and prosocial interaction are summarized in Figure 5B. First, males had lower empathy-related scores than females, regardless of the genotype (M vs F, Mann-Whitney U test: z=-3.33, *p=0.0009*; Figure 5B). Second, α5KO display lower empathy-like scores compared to their sex-matched WT controls. Finally, while α5KI male mice show no deficit as compared to WT (in contrast to α5KOs), female α5KI displayed the higher empathy-like score than any other group, including male α5KI mice and high performing female WT (Figure 5B). These observations are supported by analyses of scores yielding significant main effect of genotype and sex differences (Kruskal-Wallis test, M + F: H(2)=11.19, *p*=0.0037; M: H(2)=4.15, *p*=0.12 ns; F: H(2)=7.96, *p*=0.018; Figure 5B).

## DISCUSSION

The present study aimed at further characterizing emotion recognition and altruistic behaviors in two strains of transgenic mice expressing either a null mutation or a SNP variant of α5*nAChRs. In humans, GWAS have linked this α5SNP to an increased risk of developing nicotine dependence, schizophrenia and possibly AUDs. We previously reported that both α5 nicotinic mutants display an increase in chronic, volitional alcohol consumption, while exhibiting an opposite emotional profile reminiscent of Type I and Type II subtypes of AUD patients ^7^. Considering the transnosographic importance of social abilities, we aimed at further investigating the social emotional profile of α5 nicotinic mutants. The present study also contributes to the characterization of sex differences in these behaviors.

We first showed that, in a task aimed at distinguishing a stressed unfamiliar conspecific from a neutral one (ASDT), female WT mice were better emotional discriminators than males, exhibiting a preferential approach toward the stressed demonstrator twice superior to the males. Females were also unanimous discriminators, while a quarter of males were not. These results are in line with previous studies suggesting that female mice may have a better ability to detect negative emotions in conspecifics ^61,62^. In contrast to sex-matched WT however, neither male nor female α5KO mice did preferentially approach a stressed unfamiliar conspecific compared to a neutral one, thus revealing a deficit in their ability to perceive negative states in peers. As compared to null mutants, α5KI mice carrying the α5SNP (*rs1686699)* were less impacted in the ASDT, although they were less efficient than sex-matched WT. Importantly, female α5KI also displayed higher ASDT performances than α5KI males.

Emotion recognition in mice has been shown to rely on activation of somatostatin-expressing (SOM) interneurons of the PFC ^43^ and on oxytocinergic projections from the paraventricular nucleus of the hypothalamus to the amygdala ^39,43^. Interestingly, α5-nAChRs expressed in Vasoactive intestinal peptide-expressing (VIP) interneurons exert inhibitory control on SOM interneurons in the PFC ^50^, and in oxytocin-containing magnocellular neurons of the hypothalamic paraventricular nucleus ^63^. Moreover, blocking oxytocin receptors in the PFC of rats interfered with the pro-social motivation to free a trapped conspecific ^64^. Thus, behavioral changes observed in α5KO and α5KI mice could in part be due to differential dysregulations of the control exerted by α5*nAChR-containing interneurons on SOM interneurons, and/or to differential dysregulation of hypothalamic α5*nAChR-containing neurons which release oxytocin, a prominent part of the biological mechanism underlying empathy-related and pro-social behaviors ^28,65^. Interestingly, the ventral hippocampus also is involved in affective empathy as assessed using observational fear between cagemates ^66,67^

Regarding the prosocial act of rescuing conspecifics from the situation of spatial constraint (RTT), we first observed that WT females freed a sex-matched cagemate from constriction tubes while most of males did not. Males were more interested in alternative tubes (empty or containing white chocolate) than ones containing a trapped cagemate, and were less interested in interacting with the freed cagemate during the post-test phase, or did so antagonistically. This is consistent with previous reports showing that female rats are more motivated to free their trapped cagemates ^45^. Indeed, it has been suggested that in many species, including humans, females show more evidence of empathy-driven behaviors than males ^68–70^. However, in another study up to seventy percent of male mice were found to open the tubes restraining their sex-matched cagemate ^45,49^ and, in contrast to the present data, subjects did not open alternative empty tubes. One explanation for these differences could be the mouse strain, as we used C57BL6/J mice and not C57BL6/N as in Ueno et al. (2019) ^49^. A recent study demonstrated that mice from these two sub strains exhibit behavioral and physiological differences ^71^, that may also include empathy-related behaviors underlying this task. Another explanation could be the age of male subjects, as ours were 24 weeks old (mature adult) instead of 10 weeks old (young adults). In humans, older adults tend to exhibit more emotional empathy and pro-social behavior than younger adults ^72,73^. However, age-related effects on empathy-related behaviors have not been investigated in rodents yet. Also, we did not observe any chocolate sharing-like behavior in both male and female WT mice; but as Bartal et al. (2011) ^45^ found that rats shared chocolate chips with their freed cagemate in only half of all trials, this is likely due to the fact that we performed only one chocolate vs cagemate trial. More trials would be needed to verify whether mice also expressed chocolate sharing-like behavior.

In the RTT test, unlike WT mice, α5KO mice showed a reduced prosocial response which could be interpreted as a decrease in motivation to help their distressed peers. Similarly to sex-matched controls, the majority of α5KO males did not open the tube restraining their cagemates and primarily opened the alternative tubes including chocolate-containing and empty ones. However, when it came to the post-test social interaction, α5KO males differed from sex-matched controls by exerting high levels of aggression, which in humans is associated with deficient affective empathy ^32^. In light of recent studies showing aggression self-administration in male mice ^74^, it is possible that the opening behavior of some α5KO males was motivated by aggression seeking. Such aggression increase was not observed in α5KO females who rather showed an increase in cagemate freeing. But in contrast to sex-matched controls, α5KO females spent more time around the chocolate-containing tube than the cagemate-containing one and did not display interactive behavior toward the freed cagemate. The latter parameters coupled with their previous inability to discriminate negative from neutral emotional states in conspecifics suggest that α5KO females were more likely to free their cagemate because they enjoy the task of tube opening ^75^ rather than because they were motivated by rescuing the distressed cagemate. Taken together these results demonstrate that α5KO mice showed a decrease in empathy-related behaviors, which support their striking resemblance to the Type II AUD profile (Tochon et al. 2024). In particular, α5KO mice’s deficits in discrimination between two demonstrators in different affective states evoked the social detachment trait allocated to the Type II AUD profile ^14^. Furthermore, the aggressive behavior of α5KO males further fits the impulsive, sensation seeking profile also associated with the display of aggressive behaviors ^14,16,31^.

Female α5KI mice differed from both WT and α5KO females by displaying more interactive behavior toward the freed cagemate and by overwhelmingly opening more cagemate-containing tubes than empty ones, demonstrating a higher motivation for cagemate rescuing-like than for the tube opening act. It is worth noting that the major difference between the two social procedures was the affiliation between the subject and the non-subject mice: in the RTT, the distressed mouse to rescue was a littermate while the demonstrators to discriminate in the ASDT were unfamiliar mice. A previous study showed that male α5KI mice display a reduction of approach of a novel unfamiliar mouse compared to controls in a three-chamber sociability test ^50^. Moreover, our lab demonstrated that α5KI mice are hyper-anxious ^7^. It was previously reported that, in mice and human subjects, empathy for strangers is blunted by social stress, and blocking glucocorticoid synthesis or glucocorticoid or mineralocorticoid receptors enhances empathy in the same situation ^76^. It is therefore possible that the hyper-anxious and novelty-avoidant phenotype of α5KI mice make them perceive the stressed unfamiliar conspecific as a threat, which could hamper their exploration and explain why they were found to be bad affective state discriminators in the ASDT. The score for empathy-like behavior which takes cagemate prioritization and social-contact positivity into account during the RTT enabled us to get a good overview of the interaction of sex and mutation for the display of pro-social behavior across groups. While null mutants scored negatively in both sexes, female α5KI scored positively compared to controls making them the group with the most display of empathic-like behavior. Such increase in α5KI females particularly evoked the ‘eagerness to help others’ trait allocated to the Type I AUD profile ^14^, which further support their striking resemblance to this particular subtype.

The empathic motivation to perform rescuing-like behavior in laboratory rodents is still widely debated ^34,42,75,77–83^. The current work support that the opening behavior must be carefully interpreted. We suggest that a social release paradigm like the RTT must be coupled with at least one other task assessing the ability to recognize a distress state which is a prerequisite to express pro-social helping behavior. Analyzing the post-test interaction between the subject mouse and the trapped mouse is also essential because an opening behavior coupled with an aggression or a complete disinterest toward the distressed cagemate as observed in α5KO cannot be categorized as empathic-like or pro-social. However, an opening behavior followed by a positive interaction with the freed cagemate may either support the seeking of social contact as a rescuing motivation ^84,85^ or reveal an interest toward a distressed cagemate (and maybe some premises of consolation-like behavior). Thus, studying the relative interest for alternative tubes is also required. Prioritization of cagemate tube opening over alternative tubes as observed in α5KI females may reflect either the desire to rescue a peer or the desire to stop their stress signals ^86^. However, neither the act of opening alternative tubes, nor the act of entering it can deny the existence of empathic-like concerns. Indeed, the subject mice first learnt to open trap tubes by being themselves restrained in them. During the next trials they later had to understand that cagemates trapped inside the same device could not free themselves. Thus, the act of opening and entering alternative tubes may reflect imitative behavior, while waiting for the cagemate to free itself, rather than a higher interest in the restraint tool as proposed by Ueno, 2019 ^75^. In support, these behaviors were mostly observed in cagemate vs empty and not in chocolate vs empty trials. We point out that establishing an empathic-like score with several variables collected in the social release paradigm (opening behavior, cagemate prioritization over alternative tubes and positivity of post-test interaction) provide many analogies with human data like the sex-dependent profile and the genotype-dependent changes in nicotinic mutant strains.

Overall, we demonstrate the critical role of α5 nicotinic receptors in emotion recognition and altruistic behaviors in response to congeneric stress. We also reveal important effects of sex in WT animals and their strong modulatory impact on the phenotypes of α5 mutants. The results of our study further suggest that genetic mutations targeting a5 receptors could be a common cause of the socio-emotional deficits observed in various pathological conditions, such as addictive behaviours and schizophrenia. Future research should investigate the functional consequences of α5*nAChR mutations at the neural network level.

## Acknowledgements

We thank Uwe Maskos (Institut Pasteur, Paris) for providing us with the initial mouse founders for the local breeding of α5KO and α5KI (α5SNP) mice. This work was supported by the Centre National de la Recherche Scientifique CNRS UMR 5287 and Bordeaux University, Agence Nationale de la Recherche (ANR), French National Cancer Institute (INCa/IResP/SPA 2021-002 and INCa/ 2024-198 grants). The laboratory of VD was part of the Bordeaux Neurocampus and a member of the Laboratory of Excellence network (LabEx BRAIN), and now of the Great Research Project (GPR) Plan. As such this work was supported by French state funds managed by the Agence Nationale pour la Recherche (ANR) within the Investissements d’Avenir program under reference ANR-11IDEX-0004-02.

## Conflicts of Interest

The authors declare no competing interests.

## Author Contributions

Conceptualization, V.D. and L.T.; Methodology, V. D., N.H., C.P. and L.T.; Data Analysis, C.P. and L.T.; Writing Original Draft Preparation, C.P. and L.T.; Review & Editing, all authors; Supervision, V.D. and J.L.G.; Project Administration, V.D.; Funding Acquisition, V.D. All authors have read and agreed to the published version of the manuscript.

---

**Video 1. A ‘discriminator’ mouse in the ASDT.** This recording shows a female WT mouse displaying a preferential approach toward the stressed demonstrator (SD, left cylinder) compared to the neutral demonstrator (ND, right cylinder) during the test trial (day 4) of the ASDT. SD preference = 52.

**Video 2. A ‘non-discriminator’ mouse in the ASDT.** This recording shows a male α5KO mouse equally investigating the stressed demonstrator (SD, right cylinder) and the neutral one (ND, left cylinder) during the test trial (day 4) of the ASDT. SD preference score = −7.

**Video 3. A successful cagemate rescuing-like event in the RTT.** This recording shows a WT female mouse opening the paper-sealed tube restraining its sex-matched cagemate during the trial 11 (day 4) of the RTT.

## Notes

### Competing Interest Statement

The authors have declared no competing interest.

